# Genetic determinants of risk and survival in pulmonary arterial hypertension

**DOI:** 10.1101/317164

**Authors:** Christopher J. Rhodes, Ken Batai, Marta Bleda, Matthias Haimel, Laura Southgate, Marine Germain, Michael W. Pauciulo, Charaka Hadinnapola, Jurjan Aman, Barbara Girerd, Amit Arora, Jo Knight, Ken B. Hanscombe, Jason H. Karnes, Marika Kaakinen, Henning Gall, Anna Ulrich, Lars Harbaum, Inês Cebola, Jorge Ferrer, The NIHR BioResource – Rare Diseases Consortium UK PAH Cohort Study Consortium and the US PAH Biobank Consortium,, Ferhaan Ahmad, Philippe Amouyel, Archer Stephen L., Rahul Argula, Austin Eric D., David Badesch, Sahil Bakshi, Christopher F. Barnett, Raymond Benza, Nitin Bhatt, Harm J. Bogaard, Charles D. Burger, Murali M. Chakinala, Colin Church, John G. Coghlan, Robin Condliffe, Paul A. Corris, Cesare Danesino, Stéphanie Debette, C. Gregory Elliott, Jean Elwing, Melanie Eyries, Terry Fortin, Andre Franke, Robert P. Frantz, Adaani Frost, Joe G.N. Garcia, Stefano Ghio, Hossein-Ardeschir Ghofrani, J. Simon, R. Gibbs, John B. Harley, Hua He, Nicholas S. Hill, Russel Hirsch, Arjan C. Houweling, Luke S. Howard, Dunbar Ivy, David G. Kiely, James Klinger, Gabor Kovacs, Tim Lahm, Matthias Laudes, Katie Lutz, Rajiv D. Machado, Robert V. MacKenzie Ross, Keith Marsolo, Lisa J. Martin, Shahin Moledina, David Montani, Steven D. Nathan, Michael Newnham, Andrea Olschewski, Horst Olschewski, Ronald J. Oudiz, Willem H. Ouwehand, Andrew J. Peacock, Joanna Pepke-Zaba, Zia Rehman, Ivan M. Robbins, Dan M. Roden, Erika B. Rosenzweig, Ghulam Saydain, Laura Scelsi, Robert Schilz, Werner Seeger, Christian M. Shaffer, Robert W. Simms, Marc Simon, Olivier Sitbon, Jay Suntharalingam, Emilia Swietlik, Haiyang Tang, Alexander Y. Tchourbanov, Thenappan Thenappan, Fernando Torres, Mark R. Toshner, Carmen M. Treacy, Anton Vonk Noordegraaf, Quinten Waisfisz, Anna K. Walsworth, Robert E Walter, John Wharton, R. James White, Jeffrey Wilt, Stephen J. Wort, Delphine Yung, Allan Lawrie, Marc Humbert, Florent Soubrier, David-Alexandre Trégouët, Inga Prokopenko, Richard Kittles, Stefan Gräf, William C. Nichols, Richard C. Trembath, Ankit A. Desai, Nicholas W. Morrell, Martin R. Wilkins

## Abstract

**Background:** Pulmonary arterial hypertension (PAH) is a rare disorder leading to premature death. Rare genetic variants contribute to disease etiology but the contribution of common genetic variation to disease risk and outcome remains poorly characterized.

**Methods:** We performed two separate genome-wide association studies of PAH using data across 11,744 European-ancestry individuals (including 2,085 patients), one with genotypes from 5,895 whole genome sequences and another with genotyping array data from 5,849 further samples. Cross-validation of loci reaching genome-wide significance was sought by meta-analysis. We functionally annotated associated variants and tested associations with duration of survival.

**Findings:** A locus at *HLA-DPA1/DPB1* within the class II major histocompatibility (MHC) region and a second near *SOX17* were significantly associated with PAH. The *SOX17* locus contained two independent signals associated with PAH. Functional and epigenomic data indicate that the risk variants near *SOX17* alter gene regulation via an enhancer active in endothelial cells. PAH risk variants determined haplotype-specific enhancer activity and CRISPR-inhibition of the enhancer reduced *SOX17* expression. Analysis of median survival showed that PAH patients with two copies of the *HLA-DPA1/DPB1* risk variant had a two-fold difference (>16 years versus 8 years), compared to patients homozygous for the alternative allele.

**Interpretation:** We have found that common genetic variation at loci in *HLA-DPA1/DPB1* and an enhancer near *SOX17* are associated with PAH. Impairment of Sox17 function may be more common in PAH than suggested by rare mutations in *SOX17*. Allelic variation at *HLA-DPB1* stratifies PAH patients for survival following diagnosis, with implications for future therapeutic trial design.

**Funding:** UK NIHR, BHF, UK MRC, Dinosaur Trust, NIH/NHLBI, ERS, EMBO, Wellcome Trust, EU, AHA, ACClinPharm, Netherlands CVRI, Dutch Heart Foundation, Dutch Federation of UMC, Netherlands OHRD and RNAS, German DFG, German BMBF, APH Paris, Inserm, Université Paris-Sud, and French ANR.

## Introduction

Pulmonary arterial hypertension (PAH) refers to an uncommon but devastating disorder characterized by obliterative pulmonary vascular remodelling, leading to a progressive increase in pulmonary vascular resistance and right heart failure. The annual mortality rate for idiopathic (IPAH) and heritable PAH (HPAH) remains around 10%, despite the use of modern therapies^1,2^. In part this reflects the limited impact of licensed treatments upon the underlying pulmonary vascular pathology, which includes vascular smooth muscle and fibroblast hyperplasia, endothelial cell proliferation and inflammation^3^. Substantial variation between patients in their response to available treatments highlights underlying and inadequately characterized heterogeneity in the etiology of PAH.

Recent gene sequencing studies have revealed rare mutations in a number of genes including bone morphogenetic protein type II receptor *(BMPR2)*, potassium channels, and most recently the transcription factor *SOX17^4^*. While influencing both the risk of developing PAH and survival, rare genetic variation is found in at most 25% of patients with PAH. In the majority of PAH patients the extent of genetic contribution, including that attributable to common variation, remains largely unknown^5,6^. Therefore, we tested for genome-wide association for PAH in large international cohorts and assessed the contribution of associated regions to patient outcomes. Given the rarity of PAH, we aggregated four cohorts across North America and Europe and used a two-stage, discovery and cross-validation by meta-analysis design to assess the strength of the results.

## Methods

### PAH cohorts and genotyping

PAH was defined by hemodynamic criteria according to international guidelines^2^. Unrelated individuals with IPAH and HPAH or anorexigen-associated PAH were included. Subjects with evidence of other known causes of PAH were excluded (appendix p.2-3). All enrolled individuals provided written informed consent from their respective institutions or were included as anonymous controls under the DNA databank at Vanderbilt University-BioVU (https://victr.vanderbilt.edu/pub/biovu/) opt-out policy (appendix p.2).

Four studies were used for the analyses as summarised in Figure 1: In the *UK National Institute for Health Research BioResource (NIHRBR) for Rare Diseases study*, whole-genome sequencing (WGS, Illumina, mean depth ~35X, appendix p.2) was performed in a total of 5,895 individuals of European descent, each with a rare disorder from 16 categories or their unaffected relatives and included 847 PAH cases (Tables S1 and S2, Figure S1 appendix p.24). The concept of this study was to sequence patients with rare diseases to identify genetic influences on the pathogenesis of one rare disorder using the other rare diseases as controls, assuming that distinct rare diseases are highly unlikely to share common genetic mechanisms. This assumption was tested by repeating analyses excluding each major control group (see results below and appendix p.8).

**Figure 1.**
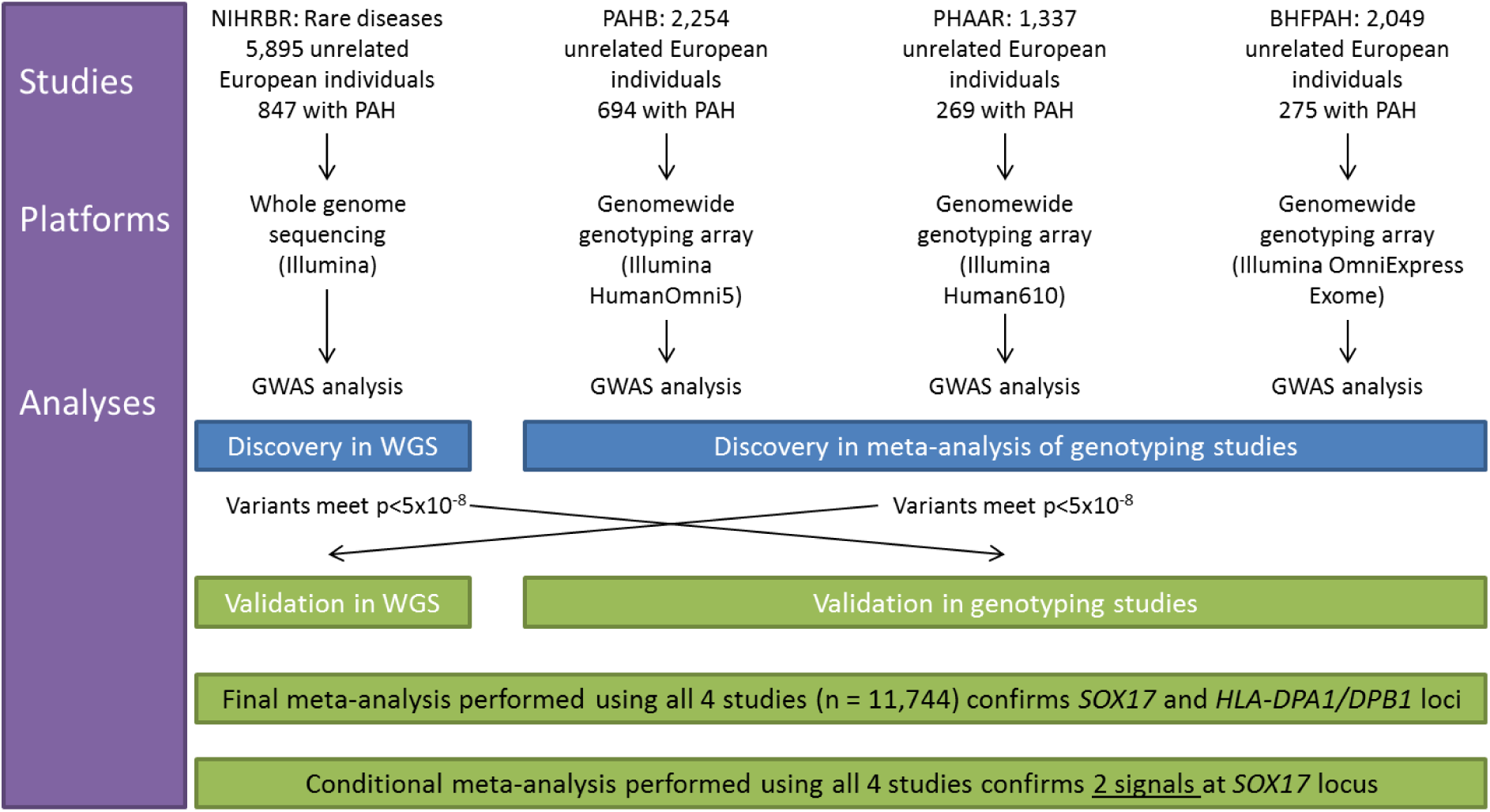
Flowchart showing study design. Four studies of European-ancestry individuals were included; one NIHRBR included rare disease patients and relatives for whom whole genome sequencing was performed. PAH patients were compared with non-PAH patients and their relatives in one discovery GWAS. Three studies, PAHB, PHAAR and BHFPAH, included PAH patients and non-PAH controls from the US, France and a mixture of European countries, respectively, for whom genome-wide array data were acquired. PAH patients were compared with non-PAH controls in each study and the results were meta-analysed in another discovery GWAS. Genome-wide significant hits from each GWAS were selected for cross-validation. Finally, all four studies were meta-analysed to provide overall associations, and conditional analysis correcting for most significant variants at each locus were used to resolve signals for multiple associations.

Three studies used genome-wide genotyping arrays: the *US National Biological Sample and Data Repository for Pulmonary Arterial Hypertension/PAH biobank (PAHB) study*, including 694 PAH cases and 1,560 controls ascertained for a large pharmacogenomic study at Vanderbilt University^7^; the *Pulmonary Hypertension Allele-Associated Risk* (PHAAR) study^5^, including 269 PAH cases and 1,068 population-based controls; and the *British Heart Foundation Pulmonary Arterial Hypertension (BHFPAH)* study, consisting of 275 PAH cases and 1,983 population-based controls, (Table S1/appendix p.12). All genotyping studies were imputed (appendix p.12) and SNPs with good imputation quality (Rsq<=0.3) taken forward for testing. Other QC steps are detailed in Table S1/appendix p.12 and appendix p.4-5.

### Association analyses

We used logistic regression to test single marker variants for genetic association with a diagnosis of PAH assuming a log-additive genetic model and adjusting for sex, read length chemistry (NIHRBR only) and for population structure using principal components analysis. Genomic inflation factor was calculated and verified to be between 1 and 1.05 for each study.

Discovery was performed in two independent sets: 1) WGS data from NIHRBR (n=5,895, including 847 PAH cases) and 2) meta-analysis of genotyping studies PAHB, PHAAR and BHFPAH (n=5,849, including 1,238 PAH cases). Cross-validation was performed and loci confirmed in a meta-analysis of all four studies using inverse-variance weighted fixed-effect meta-analyses, implemented in the GWAMA software tool^8^. We performed a conditional analysis on the lead variant in each locus to test for independent distinct signals reaching P<5×10^−8^.

LDlink was used to assess linkage disequilibrium (LD) of variants in all European populations from the 1000 Genomes Project, (https://analysistools.nci.nih.gov/LDlink/; accessed 18/07/17). Credible sets of variants considered 99% likely to include the functional causal variants were calculated by summing ranked posterior probabilities (appendix p.5&8).

### Annotation and functional assessment of the locus near *SOX17*

The locus near *SOX17* was assessed against publically available functional annotation datasets (including ENCODE and Blueprint). The locus was repressed using CRISPR-mediated inhibition in human pulmonary artery endothelial cells (hPAECs, PromoCell GmbH, Heidelberg, Germany) by transduction with a lentivirus containing a plasmid encoding the nuclease-deficient Cas9 (dCas9) fused to the repressor KRAB and a 20bp guide RNA (appendix p.6). Cells were harvested following blasticidin selection, and gene expression of *SOX17* as well as neighbouring *MRPL15* and *TMEM68* were assessed by quantitative PCR.

*In vitro* enhancer activity of the loci and variants near *SOX17* was investigated using a luciferase reporter assay. Specifically, genomic DNA (gDNA) was isolated from blood-outgrowth endothelial cells derived from a PAH patient heterozygous for the lead SNP at *SOX17* and used to clone 100bp putative enhancer regions containing the *SOX17* PAH variants. The cloned products were inserted into a luciferase reporter plasmid, which was subsequently used for transformation of stable bacteria. Picking various bacterial colonies allowed for isolation of luciferase reporter plasmids containing gDNA inserts differing only by the allele of the SNP of interest. Reporter plasmids were transfected into hPAECs by electroporation and luciferase activity was measured to quantify the enhancer function of the inserts with relevant haplotype.

### Survival analysis for lead variants and HLA alleles

All-cause mortality was used as the primary endpoint in survival analyses using Kaplan-Meier estimates and Cox regression in the *‘survival* package in R^9^. Survival was calculated from diagnosis to date of death, or censoring (NIHRBR: 31/10/16, PAHB: 01/08/17, PHAAR: 27/09/17, BHFPAH: 12/10/17), with left-truncation using date of genetic consent, and patients were censored at lung/heart-and-lung transplantation. Age and sex were included as covariates to correct for their known association with prognosis^2^.

### Analytical HLA type inference

HLA alleles and amino acids totalling 1873 features were determined by imputation from genotyped and high-quality imputed variants in the HLA region using the SNP2HLA software and the type 1 diabetes genetics consortium reference database^10^. HLA alleles and amino acids were then tested for association with the novel lead variants or case-control status by chi-squared test with FDR correction.

## Results

### Identification of PAH loci

In two separate GWAS discovery analyses, one based on a large case-control cohort that had undergone whole genome sequencing and the other comprising three genotyped case-control studies (Figure 1), we identified two loci associated with PAH reaching genome-wide significance (p<5×10^−8^, Table 1 and Figure S1/appendix p.22). One locus was within *HLA-DPA1/DPB1*, which encodes the Major Histocompatibility Complex (MHC) class II, DP alpha- and beta-chains. The second locus was 100-200kb upstream of *SOX17* which encodes the transcription factor SRY-related HMG box 17 (known as Sox17).

**Table 1.** Novel loci associated with PAH in sequenced and genotyped cohorts. Odds ratios are for association between effect allele and PAH. gnomAD is the Genome Aggregation Database, which provides information including allele frequencies in different populations.

| Variant | Chromosome and position, hg19 : Effect/ Non-effect alleles | Effect allele frequency in non-Finnish Europeans in gnomAD | Effect allele frequency in NIHRBR controls | UK NIHRBR whole genome sequencing study (n=847 cases v 5,048 controls) |  | Effect allele frequency in genotyping controls | Meta-analysis of genotyping studies US PAHB, Paris PHAAR and London BHFP AH (n= 1,238 cases v 4,611 controls) |  | Meta-analysis of all cohorts (n=2085 cases, 9659 controls, effective n=6648) |  |
| --- | --- | --- | --- | --- | --- | --- | --- | --- | --- | --- |
|  |  |  |  | Odds ratio (95% confidence intervals) | P-value |  | Odds ratio (95% confidence intervals) | P-value | Odds ratio (95% confidence intervals) | Meta-analysis P-value |
| *Lead SNPs* |  |  |  |  |  |  |  |  |  |  |
| *HLA-DPA1/DPB1*, rs2856830 | 6:33041734:C/T | 0.12 | 0.12 | 1.71 (1.48 - 1.96) | **4.41x10<sup>-14</sup>** | 0.13 | 1.44 (1.26 - 1.64) | 5.35x10<sup>-8</sup> | 1.56 (1.42 - 1.71) | **7.65x10<sup>-20</sup>** |
| *SOX17*, signal 1 rs13266183 | 8:55267612:C/T | 0.73 | 0.73 | 1.44 (1.26 - 1.64) | **4.44x10<sup>-8</sup>** | 0.74 | 1.31 (1.17 - 1.46) | 4.1x10<sup>-6</sup> | 1.36 (1.25 - 1.48) | **1.69x10<sup>-12</sup>** |
| *SOX17*, signal 2 rs10103692 | 8:55258127:G/A | 0.90 | 0.90 | 1.85 (1.47 - 2.31) | 9.51x10<sup>-8</sup> | 0.91 | 1.76 (1.45 - 2.14) | **9.84x10<sup>-9</sup>** | 1.80 (1.55 - 2.08) | **5.13x10<sup>-15</sup>** |
| *Other SNPs* |  |  |  |  |  |  |  |  |  |  |
| *HLA-DPB1* missense SNP, rs1042140 | 6:33048640:G/A | 0.23 | 0.23 | 1.38 (1.22 - 1.55) | 9.21x10<sup>-8</sup> | 0.23 | 1.44 (1.29 - 1.61) | **9.73x10<sup>-11</sup>** | 1.41 (1.30 - 1.53) | 7.13x10<sup>-17</sup> |
| *SOX17*, genotyping lead SNP rs28576721 | 8:55265980:T/C | 0.91 | 0.92 | 1.55 (1.23 - 1.95) | 1.57x10<sup>-4</sup> | 0.92 | 1.96 (1.57 - 2.43) | **1.54x10<sup>-9</sup>** | 1.75 (1.50 - 2.05) | 3.07x10<sup>-12</sup> |

### Cross-validation of PAH loci and genome-wide meta-analysis

As both the *HLA-DPA1/DPB1* and *SOX17* loci reached genome-wide significance in both discovery analyses, our cross-validation strategy simply confirmed the same alleles were more frequent in PAH in both analyses (Table 1). Next we performed genome-wide meta-analysis of all four studies, totalling 2,085 cases and 9,655 controls, which confirmed their associations with PAH (*HLA-DPA1/DPB1*, rs2856830, p=7.65×10^−20^; *SOX17*, rs10103692, p=5.13×10^−15^, Table 1 and Figure 2), and detected no further loci at genome-wide significance. Allele frequencies in the different control groups were similar between studies and to non-Finnish Europeans in the public database gnomAD (Table 1).

**Figure 2.**
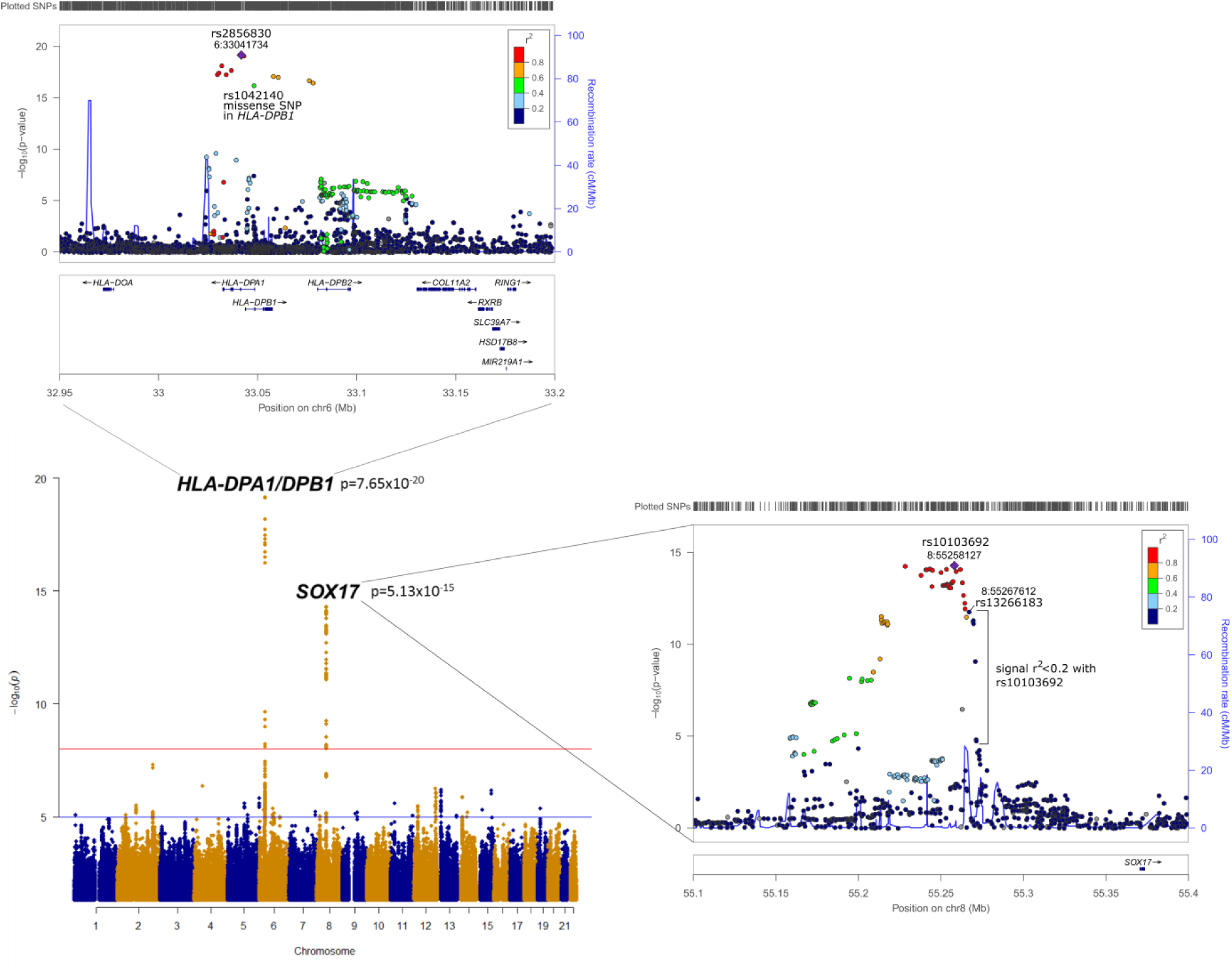
– A meta-analysis of all cohorts and regional plots of novel loci. The regional plots indicate variant location and linkage disequilibrium (LD) structure at the *HLA-DPA1/DPB1* and *SOX17* loci, respectively. At the *SOX17* locus, several variants associated with PAH are in very weak or no LD (r^2^<0.2) with the lead SNP, rs10103692. We refer to these variants as *SOX17* signal 1 and the most significant, rs13266183, is indicated. The variants coloured as in LD with rs10103692 comprise signal 2.

### Definition of key variants and independent signals within PAH loci

To determine if there was more than one signal at each locus, we performed a conditional analysis (see Methods). This confirmed that the *HLA-DPA1/DPB1* locus contained a single signal of association, but showed that the *SOX17* locus was composed of two independent signals; signal 1 is 100-103kb upstream of *SOX17* (conditional p_conditional_=9.82×10^−9^) and signal 2 is 106-200kb upstream of *SOX17* (p_conditional_=4.16×10^−11^, Figure 2 and Figure S3/appendix p.27). To narrow the variants in these loci to those 99% likely to be causal, we performed a Bayesian credible set analysis (Table S3/appendix p.14). The *HLA-DPA1/DPB1* locus included 9 SNPs (all p<9.1×10^−18^), *SOX17* signal 1 included 4 SNPs 100-103kb upstream of *SOX17* (all p<3.3×10^−8^) and *SOX17* signal 2 included 31 SNPs 106-142kb upstream of *SOX17* (all p<5.7×10^−10^).

### Testing of published loci associated with PAH and sensitivity analyses

Previous studies have reported the association of variants near *CBLN2* and *PDE1A\DNAJC10* with PAH^5,6^. These common variant signals showed no association with PAH in the combined NIHRBR, PAHB and BHFPAH cohorts (p=0.17; and p=0.24, respectively; Table S2/appendix p.13). Sensitivity analyses excluding pathogenic *BMPR2* variant carriers, all pathogenic rare variant carriers or controls from different disease groups yielded similar results to the main analyses (appendix p.8).

### Functional impact of PAH locus upstream of *SOX17*

To search for evidence of regulatory elements in relevant tissues at *SOX17* signal 1 and signal 2, we examined publically available epigenomic data (including histone modifications, Figure 3 and Figure S2/appendix p.23). This identified several putative enhancer elements active in both lung tissue and endothelial cells (Figure 3). One of these (around hg19-chr8:55.270Mb) contains a cluster of three out of four credible variants from *SOX17* signal 1 (Figure 3). Another (around hg19-chr8:55.252Mb) contains 1 credible variant from *SOX17* signal 2. Of these variants, rs10958403 in signal 1 and rs765727 in signal 2 overlap a DNAse hypersensitivity signal, which indicates accessible chromatin (allowing binding of transcription factors), detected in human pulmonary artery endothelial cells (hPAECs, Figure 3).

**Figure 3.**
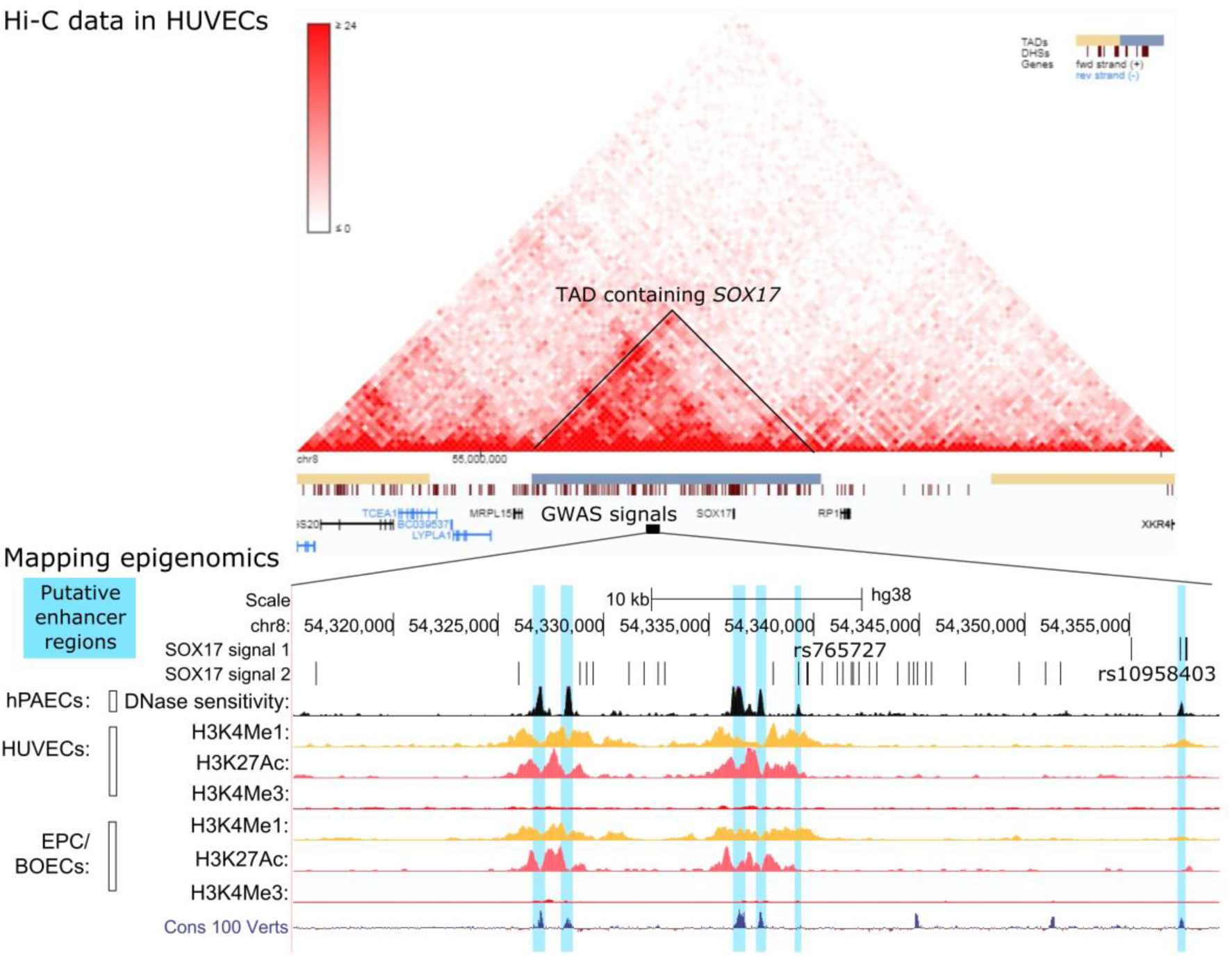
- *In silico* analysis of SOX17 locus. Hi-C data from human umbilical vein endothelial cells indicates regions of DNA found in close proximity in the 3D structure. The GWAS locus position is indicated by a black box, overlapping a TAD indicated in blue which contains only *SOX17*. Mapping of *SOX17* locus variants associated with PAH with public epigenomic data is underneath Hi-C data. The credible set indicates positions of variants 99% likely to contain the causal variants. Transcription factor binding sites as determined by ChIP-Seq experiments of 161 factors from ENCODE with Factorbook Motifs are shown; H indicates binding site in HeLa-S3 cervix adenocarcinoma cells, U indicates binding site in human umbilical vein endothelial cells (HUVEC). Auxiliary hidden markov models (HMM), which summarize epigenomic data to predict the functional status of genomic regions in different tissues/cells, are shown. Epigenomic data in endothelial cells (EC) including HUVEC, human pulmonary artery ECs (hPAECs) and endothelial progenitor cells (EPC), also known as blood outgrowth ECs (BOEC), indicate areas likely to contain active regulatory regions and promoters. Markers include histone 3 lysine 4 monomethylation (H3K4Me1, often found in enhancers) and trimethylation (H3K4Me3 strongly observed in promoters) and lysine 27 acetylation (often found in active regulatory regions). The blue areas indicate where epigenomic data suggest a putative enhancer region, some overlapped by variants associated with PAH. These regions were cloned for the luciferase reporter experiments (results in Figure 4B).

To study the effects of the PAH risk variants on the putative enhancers defined by the epigenomic signals, we developed reporter constructs containing 100bp of the regions containing either the risk allele or non-risk alleles at each of the 4 SNPs using genomic DNA from a patient heterozygous for both *SOX17* signals. A haplotype-specific reporter assay in hPAECs confirmed that the regions containing either rs10958403 or rs765727 exhibited enhancer activity (between 3 and 6-fold induction of luciferase/Renilla ratio, *p*<0.001), whereas constructs containing rs12674755 or rs12677277 had no effect compared to the empty vector control. We also observed haplotype-specific activity with the active constructs, which differed only by the alleles at PAH-associated risk variants rs10958403 or rs765727 (both p<0.05, Figure 4B).

**Figure 4.**
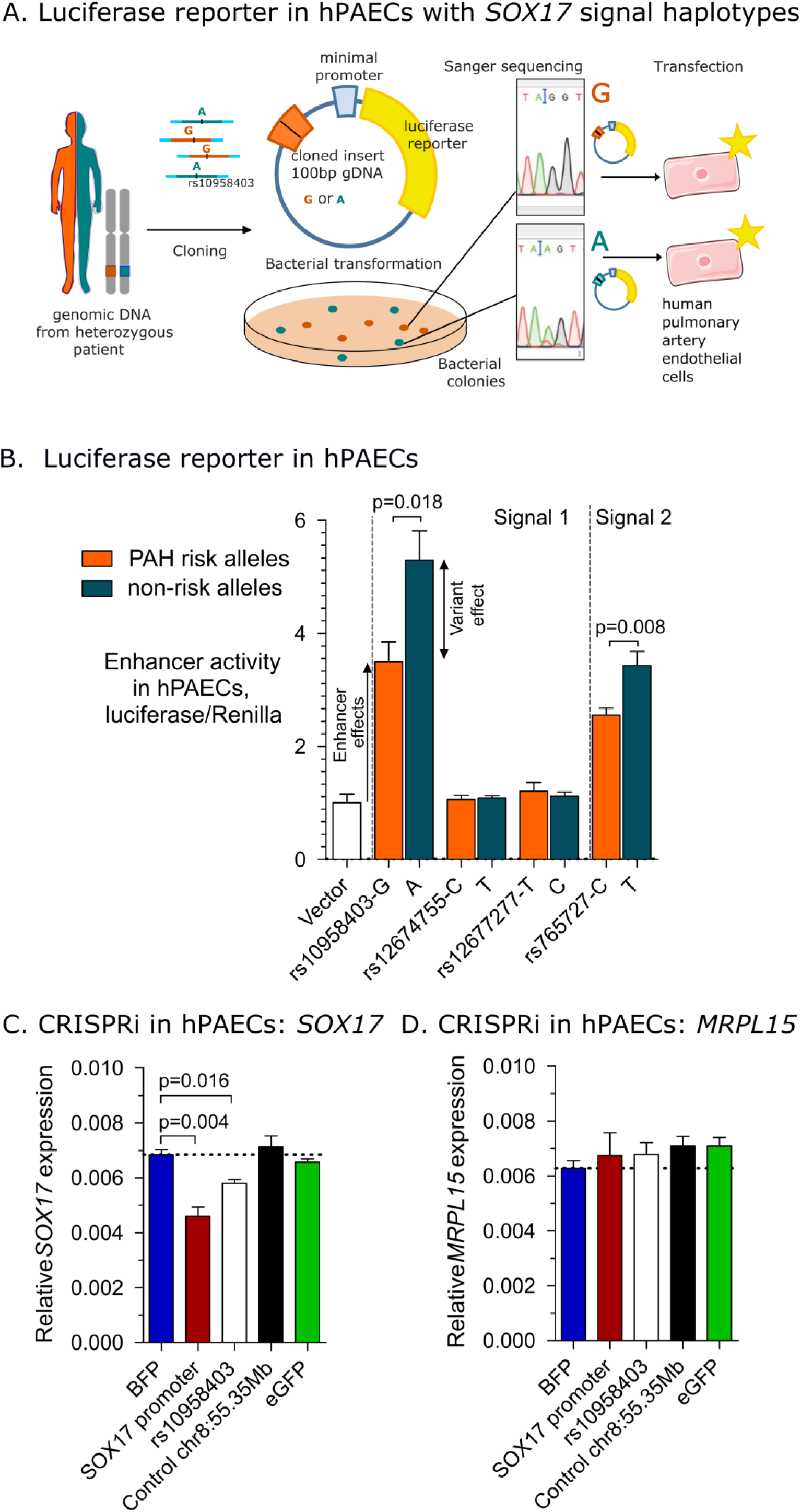
**A**. Cartoon describing process for haplotype specific reporter construct derivation. 100bp gDNA inserts containing *SOX17* SNPs are isolated from blood outgrowth endothelial cells derived from a PAH patient heterozygous for the *SOX17* SNPs. Colonies of transformed bacteria can be sequenced to determine allele present in product. Transfection of luciferase reporter constructs containing inserts into human pulmonary artery endothelial cells allows for determination of luciferase activity. **B**. Luciferase reporter assay results. Luciferase/Renilla ratios relative to the empty vector demonstrate haplotype-dependent enhancement of promoter activity. Enhancer effects were tested by one way analysis of variance followed by Dunnett’s post-hoc tests - rs10958403-G/A and rs765727-C/T were both p<0.0001 significant versus empty vector, variant effects of these 2 SNPs were tested by t-test. Mean±SEM of n=5 experiments. **C**. Relative expression of *Sox17:beta-actin* in hPAECs upon CRISPR-mediated repression of the near *SOX17* GWAS locus. Mean±SEM of n=4 measurements in a representative experiment. 3 further experiments showed consistent results. BFP, blue fluorescent protein; eGFP, enhanced green fluorescent protein; and control, which refers to a region between the enhancer region and the *SOX17* gene that is negative for regulatory markers, are used as negative controls. The *SOX17* promoter was targeted as a positive control of repression. Significance shown vs. BFP by Dunnett’s post-hoc analysis. **D**. Relative expression of *Mrpl15:beta-actin* in hPAECs upon CRISPR-mediated repression of the GWAS locus.

DNA folding patterns determined by Hi-C data from lung tissue and endothelial cells (human umbilical vein endothelial cells, human microvascular endothelial cells, Figure 3A) indicate that the *SOX17* PAH locus resides in a defined topologically associated domain (TAD) in which the only gene found, and thus likely target of any regulatory elements in this region, is *SOX17*. To test this, we performed CRISPR-mediated inhibition of the *SOX17* signal 1 region in hPAECs. This resulted in selective down-regulation of *SOX17* expression but not the expression of neighbouring *MRPL15* and *TMEM68* genes, suggesting that the enhancers in this locus specifically regulate *SOX17* (Figure 4C-D and Figure S3/appendix p.25).

### Associations of PAH loci with clinical outcomes

We investigated whether the *HLA-DPA1/DPB1* and *SOX17* variants influence clinical outcomes in PAH, specifically all-cause mortality. The *HLA-DPA1/DPB1* rs2856830 genotype, but not the *SOX17* locus, was strongly associated with survival (Figure 5). Median survival from diagnosis in the NIHRBR and PAHB patients with the C/C homozygous genotype was double (median[95%CI] =16.34[12.34->16.34] years) that of the T/T genotype (median[95%CI] = 8.05[5.76-11.3] years). Cox regression survival analyses showed that the rs2856830 T/T genotype conferred an increased annual risk of death in PAH of 97% (Figure 5B).

**Figure 5.**
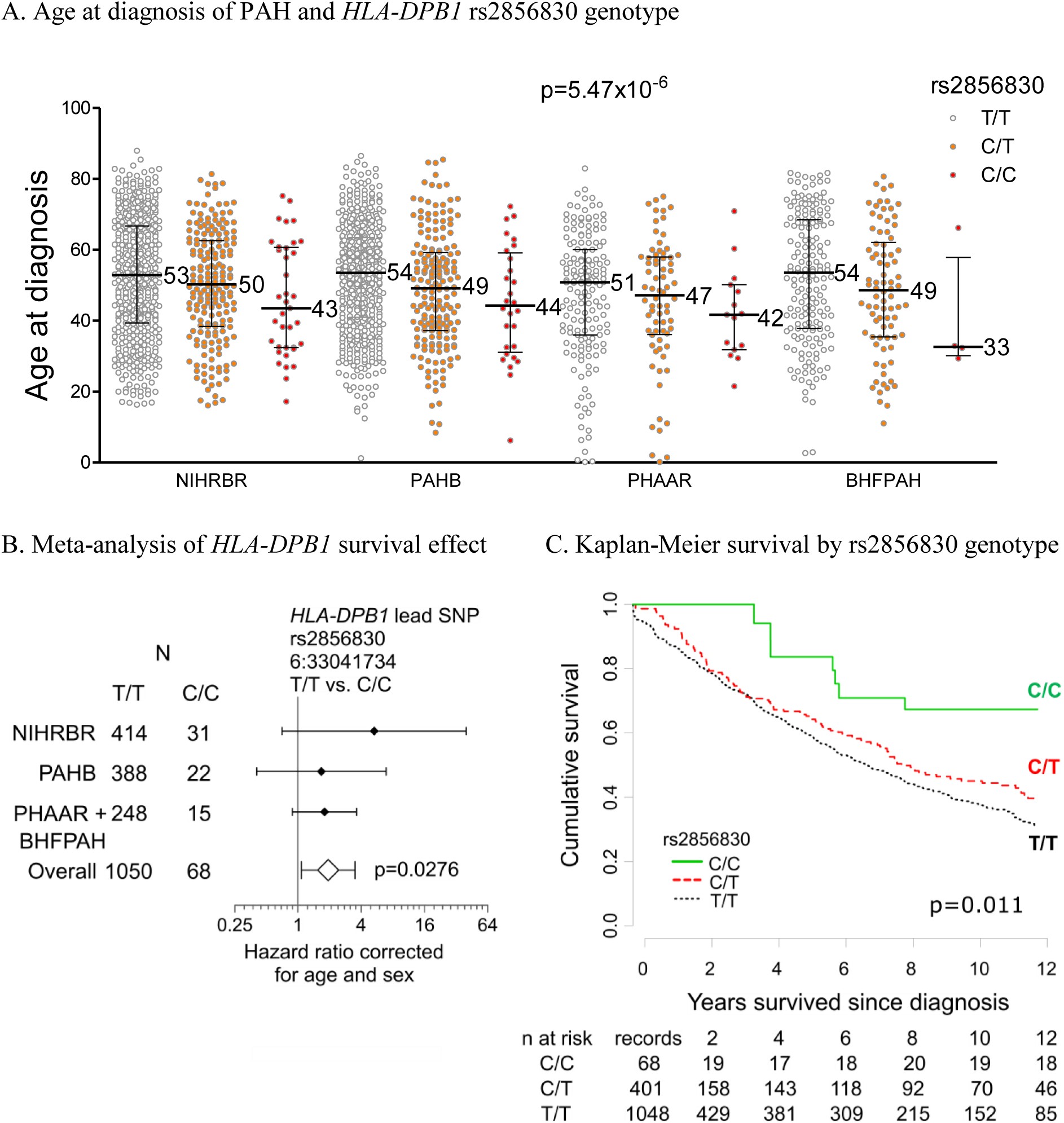
– Clinical impact of *HLA-DPB1* rs2856830. **A**. Age at diagnosis by genotype in four cohorts of PAH patients. Bars indicate medians and interquartile range, numbers given are median values in subgroups. p-value shown is from linear regression model correcting for cohort differences. **B**. Forest plot showing hazard ratios for the rs2856830T/T vs C/C genotypes, corrected for age and sex in Cox regression survival analyses in each PAH cohort, individually and with meta-analysis results. **C**. Kaplan-Meier survival plot in PAH patients divided into groups based on the genotype of *HLA-DPA1/DPB1* SNP rs2856830 in all cohorts. N at risk indicates numbers at risk in each time period, which increases as truncated patients are recruited into the study after diagnosis and decreases as patient follow-up ends. Significance from log rank test is given.

Sensitivity analyses excluding pathogenic *BMPR2* variant carriers, all pathogenic rare variant carriers and patients diagnosed in previous decades who may have been exposed to different treatment regimens gave results similar to the main analyses (appendix p.8).

We tested both loci for association with other clinical variables, including disease severity measures and comorbidities (Tables S5 and S6/appendix p.16-17). The C allele at *HLA-DPA1/DPB1* lead SNP rs2856830 was associated with younger age at diagnosis (Figure 5A), with C/C homozygotes presenting a decade earlier (Table S5/appendix p.16). The rs2856830 genotype was not associated with vasoresponder status.

### PAH locus at *HLA-DPA1/DPB1*

The *HLA-DPA1/DPB1* locus included a missense variant rs1042140 in *HLA-DPB1* reaching genome-wide significance, (Table 1) in partial LD (r^2^=0.45 with lead rs2856830 in Europeans). The SNP, rs1042140, determines a glutamic acid (Glu^69^) or a lysine at amino acid residue 69. To determine specific HLA alleles associated with the lead variant, rs2856830, we imputed HLA types from the genotype data. These types are represented by digit codes, where the first 2 digits represent related groups of similar alleles (e.g. *DPB1*\*02), and 4 digits represent specific proteins with distinct amino acid sequences (e.g. *DPB1*\*02:01). We found that the PAH-enriched C allele of rs2856830 was associated with HLA-*DPB1*\*02:01/02:02/16:01 (all p<1 × 10^−9^ after FDR correction, Table 2 and Table S7/appendix p. 18), which all contain the Glu^69^ residue. The most numerous *DPB1*\*02:01 and *DPB1*\*04:01 alleles were associated with survival in PAH patients (hazard ratio, HR[95%CI]=0.70[0.49-1.00] and HR[95%CI]=1.33[1.04-1.70], respectively, Table 2).

**Table 2.**
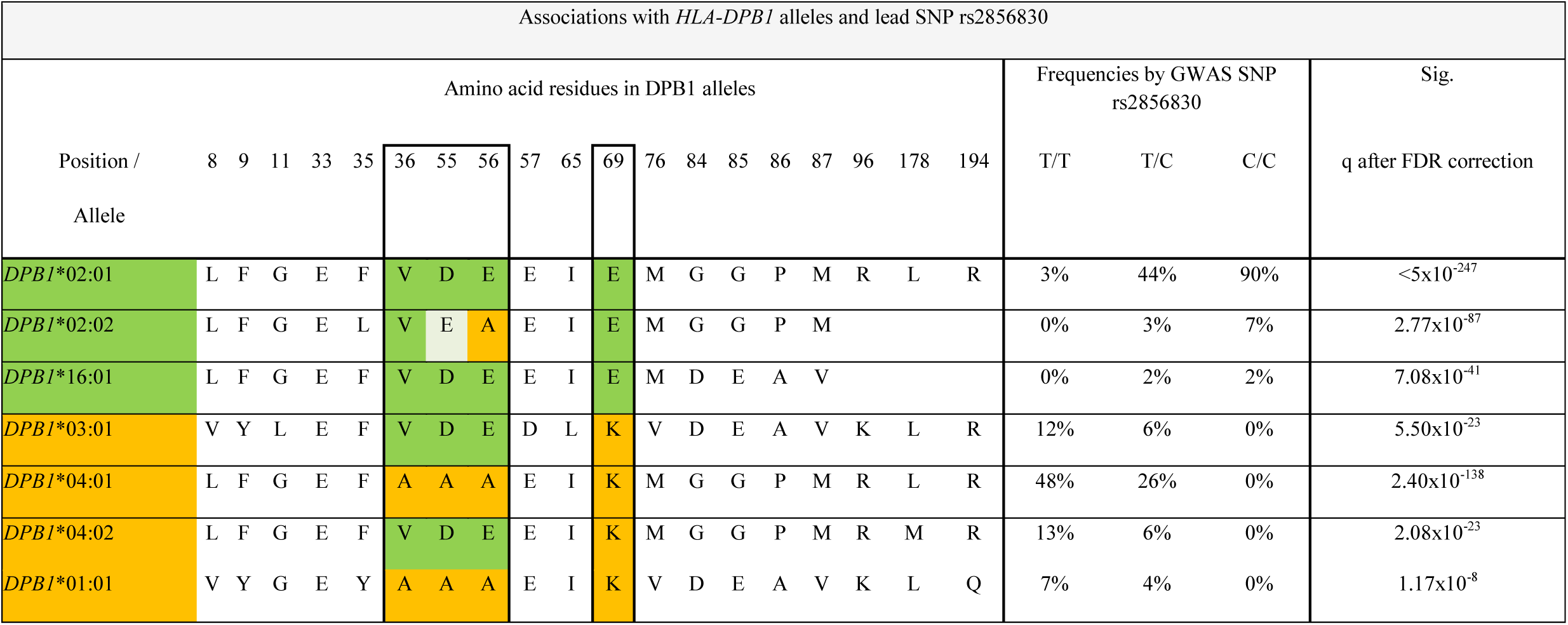
Associations of *HLA-DPB1* alleles with the lead SNP rs2856830. Orange indicates alleles and residues depleted in PAH cases and green indicates those enriched in PAH cases.

### Frequency of PAH risk alleles

The risk alleles at both signals within the *SOX17* locus are common (risk allele frequencies are rs13266183-C=74% and rs9298503-C=92%, respectively), such that 59% of PAH cases were homozygous for the risk allele at both *SOX17* SNPs, compared to only 46% of controls.

The alleles at *HLA-DPB1* associated with the poorest outcomes are also common (risk allele frequency of rs2856830-T=86%), such that 69% of PAH patients had the T/T genotype associated with the poorest outcomes and 95% had at least one T allele.

## Discussion

Through a meta-analysis of 11,744 individuals we have established loci at *HLA-DPA1/DPB1* and at an enhancer upstream of *SOX17* associated with PAH disease risk. Polymorphic variation at the *HLA-DPA1/DPB1* locus is strongly associated with both the age at diagnosis and prognosis in PAH. Common genetic variants in the enhancer region of *SOX17* are biologically plausible candidates for susceptibility to pulmonary vascular disease.

Both *in silico* and experimental analyses of the common variants upstream of the *SOX17* gene suggest they influence susceptibility to PAH through regulation of *SOX17* expression. We have recently reported enrichment and familial segregation in PAH of causal rare deleterious variation in *SOX17*, implicating this gene in the pathogenesis of PAH^4^. Sox17 is involved in the development of the endoderm^11–13^, vascular endothelium, haematopoietic cells^14^ and cardiomyocytes^15,16^. Sox17 also determines the endothelial fate of CD34+ progenitor cells de-differentiated from fibroblasts^17^. Deletion in the mouse leads to abnormal pulmonary vascular development, poor distal lung perfusion and biventricular hypertrophy^18^. Sox17 is a pro-angiogenic transcription factor and interacts with well-established endothelial molecular mediators^19,20^; reduction of Sox17 in endothelial cells through Notch activation (itself associated with BMPR2 signalling^21^) restricts angiogenesis^19^. Conversely, vascular endothelial growth factor (VEGF) upregulates Sox17 and, as part of a positive feedback loop, Sox17 promotes expression of VEGF receptor 2 (VEGFR2)^20^. This is relevant as inhibition of VEGFR2 results in severe pulmonary hypertension in established preclinical models^22^.

We report *HLA-DPB1* alleles associate with PAH and have a pivotal role in determining disease progression. The beneficial effect of the C/C genotype at rs2856830 on survival is greater than that of any current PAH-specific drug treatment^23^, with the exception of calcium channel blockers which are effective in a small subgroup (less than 10%) of PAH patients classified as “vasoresponders”^2^. Patients with the C allele at rs2856830 presented at a significantly younger age, but the association of the *HLA-DPB1* SNP with survival remains significant after correction for both age and sex. Clinical HLA typing or rs2856830 genotyping could improve risk stratification both in clinical practice and in clinical trials, where over-representation of the C/C genotype in one treatment arm could significantly impact outcomes.

The mechanism of rs2856830 involvement in PAH is likely through its association with specific *HLA-DPB1* alleles. Class II *(HLA-DRB1, -DQB1*, and *-DPB1)* antigen-presenting proteins play critical roles in the adaptive immune response^24,25^. The *HLA-DPB1* alleles associated with rs2856830 (*HLA-*DPB1*02:01/02:02/16:01) in the current study have also previously been linked to susceptibility to hard metal lung diseases such as berylliosis^26,27^. A number of individual amino acid residues in the peptide-binding pockets of the *HLA-DPB1* molecule influence its function and T-cell recognition, either by changing peptide antigen binding or the conformation of the peptide-binding groove^28^. *HLA-*DPB1*02:01/02:02/16:01 all contain a glutamate at position 69 and a valine at position 36 that reduce the risk of clinical deterioration. These same residues are essential for T-cell activation and cytokine production in berylliosis^29,30^. The potential role of this modification in antigen binding, autoimmune response and vascular damage in PAH demands further investigation.

We have shown in a rare disorder that common variation can drive significant clinical differences in presentation and outcomes. Furthermore, a common non-coding variant can regulate expression of a gene linked by rare, deleterious mutations to the same pathology. *HLA-DPB1*, and wider immune regulatory pathways, should be considered a priority for patient stratification and investigation of new treatments in PAH. *SOX17* is a key endothelial regulator and its dysfunction in PAH may be more common than heritable cases suggest.

## Acknowledgements

We gratefully acknowledge the participation of patients recruited to the UK National Institute of Health Research BioResource - Rare Diseases (NIHR BR-RD) Study. We thank the NIHR BR-RD staff and co-ordination teams at the University of Cambridge, and the research nurses and coordinators at the specialist pulmonary hypertension centres involved in this study. We are also grateful to Dr Jenny Thomson and Caroline Langman for invaluable assistance in BHFPAH patient recruitment. The UK National Cohort of Idiopathic and Heritable PAH is supported by the NIHR BR-RD, the British Heart Foundation (SP/12/12/29836), the BHF Cambridge Centre of Cardiovascular Research Excellence, and the UK Medical Research Council (MR/K020919/1), the Dinosaur Trust, BHF Programme grants to RCT (RG/08/006/25302), NWM (RG/13/4/30107) and MRW (RG/10/16/28575), and the UK National Institute for Health Research Cambridge Biomedical Research Centre. We also gratefully acknowledge the participation of patients recruited to the National Institutes of Health/National Heart, Lung, and Blood Institute-sponsored National Biological Sample and Data Repository for PAH (aka PAH Biobank). We thank the physicians, research nurses and coordinators at the 40 pulmonary hypertension centers across the US involved in the PAH Biobank. For a listing of investigators and enrolling centers, please visit www.pahbiobank.org. Vanderbilt University Medical Center’s BioVU projects are supported by numerous sources: institutional funding, private agencies, and federal grants. These include the NIH funded Shared Instrumentation Grant S10RR025141; CTSA grants UL1TR002243, UL1TR000445, and UL1RR024975. The genotyping of the VESPA samples was supported by RC2GM092618. The authors acknowledge use of BRC Core Facilities provided by the financial support from the Department of Health via the National Institute for Health Research (NIHR) comprehensive Biomedical Research Centre award to Guy’s and St Thomas’ NHS Foundation Trust in partnership with King’s College London and King’s College Hospital NHS Foundation Trust. NWM is a British Heart Foundation Professor and National Institute of Health Research (NIHR) Senior Investigator. CH is a NIHR Rare Disease Translational Research Collaboration Clinical PhD Fellow. CJR is supported by a British Heart Foundation Intermediate Basic Science Research Fellowship (FS/15/59/31839). LH and JA are the recipients of ERS and joint ERS/EMBO Long-Term Research fellowships LTRF 2016–6884 and LTRF 201701-00072. AL is supported by a British Heart Foundation Senior Basic Science Research Fellowship (FS/13/48/30453). LS is supported by the Wellcome Trust Institutional Strategic Support Fund (204809/Z/16/Z) awarded to St. George’s, University of London. IP is supported by the Wellcome Trust (WT205915), and the EU H2020 (DYNAhealth, project number 633595). Funding for the PAH Biobank is provided by NIH/NHLBI HL105333. WCN and MWP are supported by NIH NHLBI HL105333. JHK receives support from the American Heart Association (16SDG29090005) and the American College of Clinical Pharmacy Research Institute (Futures Grant). AAD receives support from NIH NHLBI R01HL136603. JF is supported by the Wellcome Trust (WT101033). JBH is supported by eMERGE U01 (NHGRI U01 HG008666). We acknowledge the support of the Imperial NIHR Clinical Research Facility and Biomedical Research Centre, the Netherlands CardioVascular Research Initiative, the Dutch Heart Foundation, Dutch Federation of University Medical Centres, the Netherlands Organisation for Health Research and Development and the Royal Netherlands Academy of Sciences. MRW and HAG receive funding from German Research Foundation (DFG) SFB1213, project A09. The popgen 2.0 network is supported by a grant from the German Ministry for Education and Research (01EY1103). We thank all the patients and their families who contributed to this research and the Pulmonary Hypertension Association (UK) for their support.

